# Pannexin-1 mediated ATP release in adipocytes is sensitive to glucose and insulin and modulates lipolysis and macrophage migration

**DOI:** 10.1101/380469

**Authors:** Marco Tozzi, Jacob B. Hansen, Ivana Novak

## Abstract

**One-sentence summary:** Insulin inhibits ATP release in adipocytes

**Abstract:** Extracellular ATP signaling is involved in many physiological and pathophysiological processes, and purinergic receptors are targets for drug therapy in several diseases, including obesity and diabetes. Adipose tissue has crucial functions in lipid and glucose metabolism and adipocytes express purinergic receptors. However, the sources of extracellular ATP in adipose tissue are not yet characterized.

Here, we show that upon adrenergic stimulation white adipocytes release ATP through the pannexin-1 pore that is regulated by a cAMP-PKA dependent pathway. The ATP release correlates with increased cell metabolism, and extracellular ATP induces Ca^2+^ signaling and lipolysis in adipocytes and promotes macrophages migration. Most importantly, ATP release is markedly inhibited by insulin, and thereby auto/paracrine purinergic signaling in adipose tissue would be attenuated. Furthermore, we define the signaling pathway for insulin regulated ATP release.

Our findings reveal the insulin-pannexin-1-purinergic signaling cross-talk in adipose tissue and we propose that deregulation of this signaling may underlie adipose tissue inflammation and type-2 diabetes.

## Introduction

Purinergic signaling forms an extensive regulatory system in many physiological and pathophysiological processes in mammalian cells. Extracellular nucleosides and nucleotides exert their action via a number of purinergic receptors (Adenosine, P2X and P2Y), which are well characterized in many cells, and provide attractive therapeutic targets in e.g. neurologic and cardiovascular diseases, inflammation, osteoporosis, cancer and diabetes (1). Our knowledge about the mechanisms leading to release of nucleotides is less settled. Nevertheless, it is known that most cells have a basal “resting” ATP release (2) and, depending on the cell type, ATP release can be also stimulated by hormones, agonists, changes in membrane voltage, mechanical and chemical stress (3). There are two main types of ATP release mechanisms. One is based on exocytosis of vesicles containing ATP, which is accumulated there by action of the vesicular nucleotide transporter (VNUT, *SLC17A9*), and is found in e.g. neurons, endocrine and exocrine cells (4). Another type of ATP release mechanism includes plasma membrane ion channels and transporters such as pannexin-1, connexin hemichannels, cystic fibrosis transmembrane conductance regulator (CFTR), cell-volume sensitive anion channels and maxi-anion channels and even the P2X7 receptors (3,5,6). Recent research indicates that the potentially most important mechanism for ATP release may be pannexin-1 (Panx 1), which is also implicated in inflammation, chronic pain, cardiovascular and other diseases (7). Panx 1 hemichannel subunits assemble in hexamers and behave as ion channels and pores, allowing passage of molecules up to 1.5 kDa. Panx 1 is activated by a number of stimuli, including intracellular calcium, P2X7 receptors, cell depolarization and hypoxia; and it is irreversibly activated by caspase 3/7 (8,9).

There are strong links between adipose tissue dysfunction and impaired glucose homeostasis reflected in obesity and type-2 diabetes (10,11), and accumulating evidence indicates that changes in purinergic signaling are associated with these pathologies (12–15). Dysregulated white adipocytes could contribute to obesity and low-level chronic inflammation of white adipose tissue characterized by macrophage infiltration (16–18). White adipose tissue function in glucose and lipid homeostasis and adipokines secretion is regulated by insulin, metabolites, and importantly, by sympathetic nerves releasing norepinephrine that act through adrenergic receptors (19,20). It is widely accepted that, together with catecholamines, sympathetic nerves also release ATP (21). Following ATP hydrolysis by ecto-nucleotidases, resulting adenosine binds to adenosine receptors that are well characterized in adipocytes (14). Adipocytes also express various P2Y and P2X receptors utilizing Ca^2+^ and cAMP signaling pathways and stimulating multiple responses, including the pro-inflammatory response via P2X7 receptors (14,22). Therefore, it is important to understand whether there are extracellular ATP sources other than sympathetic nerves in adipose tissue. One study shows that adrenergic stimulation causes the release of ATP from white adipocytes and that Panx1 might be involved (23). In brown adipose tissue, ATP is released from sympathetic nerves (24). VNUT expression was detected by immunohistochemistry in nerves and brown adipocytes, but not in white adipocytes (25). Clearly, ATP release mechanisms are not yet well established or characterized and events that lead to ATP release in adipocytes are unknown.

A better understanding of the mechanism of ATP release in adipocytes might be important for the following reasons: if the ATP release was regulated and specific, it may provide an early step for therapeutic intervention upstream of more complex and manifold adenosine and P2 receptors; it may provide an understanding of adipocyte metabolism and purinergic signaling within the adipose tissue microenvironment. Thus, the aim of our study was to elucidate the mechanism and regulation of ATP release in white adipocytes, and to evaluate the role of extracellular ATP as potential autocrine and paracrine signal. We show that upon adrenergic stimulation mature white adipocytes release ATP through Panx1 that is activated in a cAMP- and PKA-dependent manner. Furthermore, we provide evidence that extracellular ATP evokes calcium signaling and modulates lipolysis in white adipocytes and stimulates the recruitment of macrophages. Importantly, the ATP release pathway is sensitive to glucose and markedly inhibited by insulin. This novel insulin-Panx1-purinergic signaling cross talk/cooperation may provide new framework for understanding processes in adipose tissue that are deregulated in type-2 diabetes.

## Results

### Adrenergically stimulated adipocytes release ATP through Panx 1

To determine whether adipocytes are capable of releasing ATP, we used a well-established model for white adipocytes (murine cell line C3H10T1/2). Differentiated, mature adipocytes were stimulated with various agonists: α-and β-adrenergic agonists (phenylephrine and isoproterenol, respectively); adenylyl cyclase activator (forskolin); and calcium ionophore (ionomycin); and followed the extracellular ATP concentration online using live-cell luminescence (Fig. 1A). Adrenergically stimulated mature adipocytes revealed a robust and controlled ATP release, which began around 50 min and peaked about 120 min after stimulation. Forskolin induced an even higher ATP release and followed the same dynamics as adrenergic stimulation. Ionomycin had no effect on ATP release (although it did evoke calcium transients; see below), and the lack of response was similar to the control (Fig. 1A, C). Notably, pre-adipocytes (undifferentiated C3H10T1/2 cells) did not show any ATP release with adrenergic agonists, forskolin or ionomycin stimulation (Fig. 1B).

**Fig. 1.**
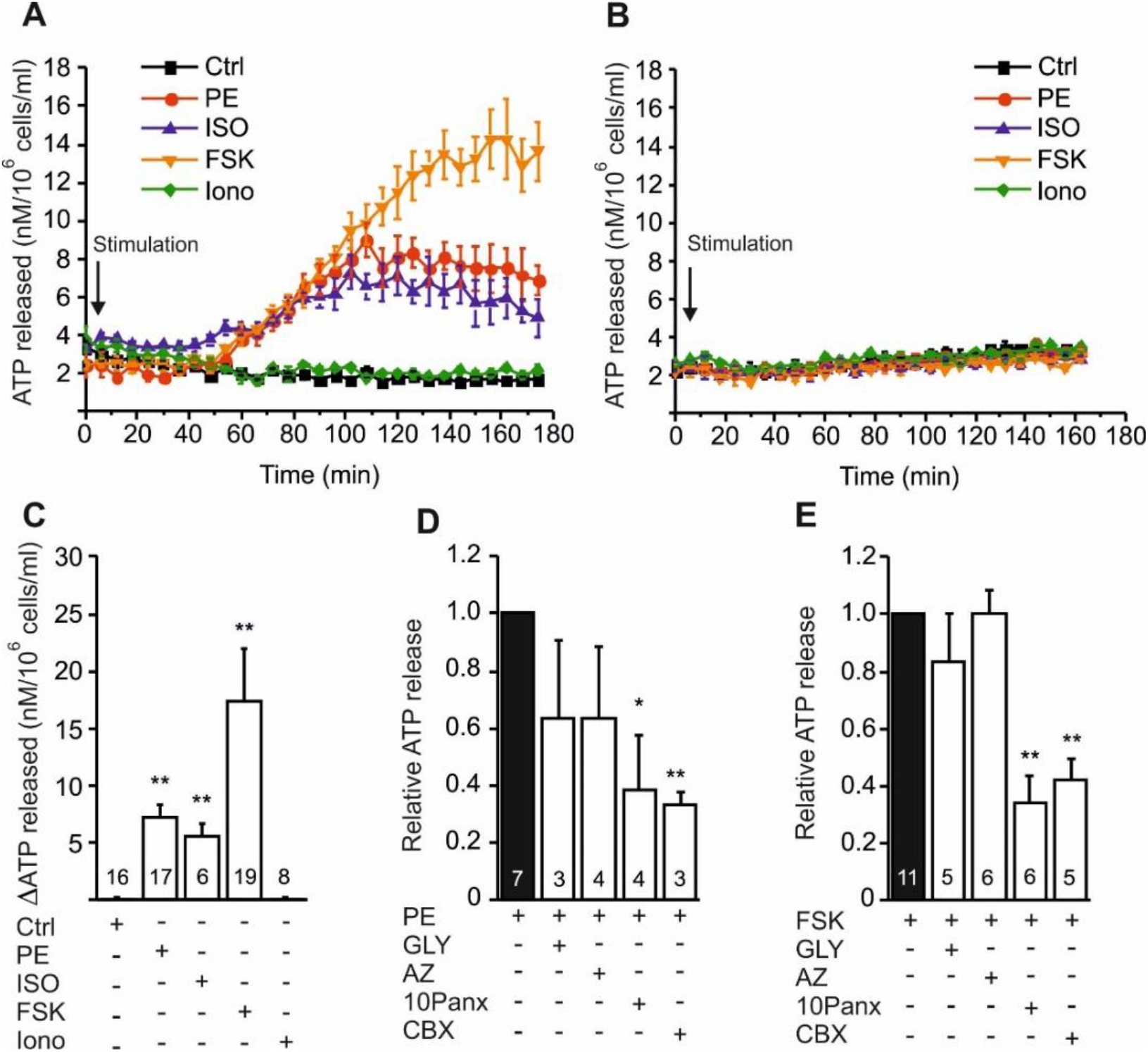
Adrenergic stimulation evokes ATP release via Panx 1 channel in white adipocytes. **(A, B)** Representative experiment showing the time-course of ATP release in differentiated (a) and undifferentiated (b) C3H10T1/2 adipocytes after stimulation with a control buffer (Ctrl), 5 μM phenylephrine (PE), 10 μM forskolin (FSK), 1 μM isoproterenol (ISO) and 1 μM ionomycin (Iono). Data represent means ± s.e.m. of 3 technical replicates. **(C)** Quantification of ATP release is in mature adipocytes is shown as a difference between the basal extracellular ATP (before stimulation) and the peak release with the indicated agonist. **(D)** Normalized phenylephrine- and (e) forskolin-induced ATP release in presence of 50 μM sodium glyoxylate (GLY), 10 μM AZ10606120 (AZ), 100 μM ^10^Panx (10Panx) and 30 μM Carbenoxolone (CBX). Data (c-e) represent means ± s.e.m., numbers denote the number of independent experiments and significant differences p<0.05 (*) and p<0.01 (**) are indicated.

The ATP release mechanism was tested using the VNUT inhibitor (glyoxylate), the P2X7 receptor inhibitor (AZ10606120), and two different Panx1 inhibitors, the mimetic inhibitory peptide (^10^Panx) and the connexin blocker (carbenoxolone) (Fig. 1D, E). The latter inhibitor was used at a relatively low concentration, which is known to be more selective for Panx1 rather than connexin hemichannels (8). Phenylephrine and forskolin induced ATP release was significantly diminished by both Panx1 inhibitors, but not by VNUT or P2X7 receptor inhibitors (Fig. 1D, E), indicating that Panx1 is the major ATP efflux pathway in adrenergically stimulated adipocytes.

### Mature adipocytes express Panx1C and form a pore

Expression of Panx1 in pre-adipocytes and mature adipocytes was determined by PCR analysis (Fig. 2A). Mature adipocytes showed two distinct PCR products, most likely corresponding to the two putative Panx1 splice variants: the well characterized NM_019482.2 (698 bp); and the predicted NM_019482 (477 bp). These variants are already annotated in the mouse genome mm9 assembly in the reference sequence database RefSeq, and here we name these as Panx1A and Panx1C, respectively, corresponding to similar variants in the rat (26). In contrast, pre-adipocytes seem to express only the longer Panx1A isoform (Fig. 2A). Western blot analysis shows that an anti-Panx1 antibody recognized the Panx1A isoform at ~48 kDa in both pre-adipocytes and mature adipocytes and a more abundant Panx1C isoform at ~45 kDa present only in mature adipocytes (Fig. 2B). Quantification of the two bands clearly shows that the expression of the ~48 kDa isoform is comparable between pre-adipocytes and mature adipocytes, while the newly characterized ~45 kDa isoform (Panx1C) is markedly upregulated in the mature state (Fig. 2C).

**Fig. 2.**
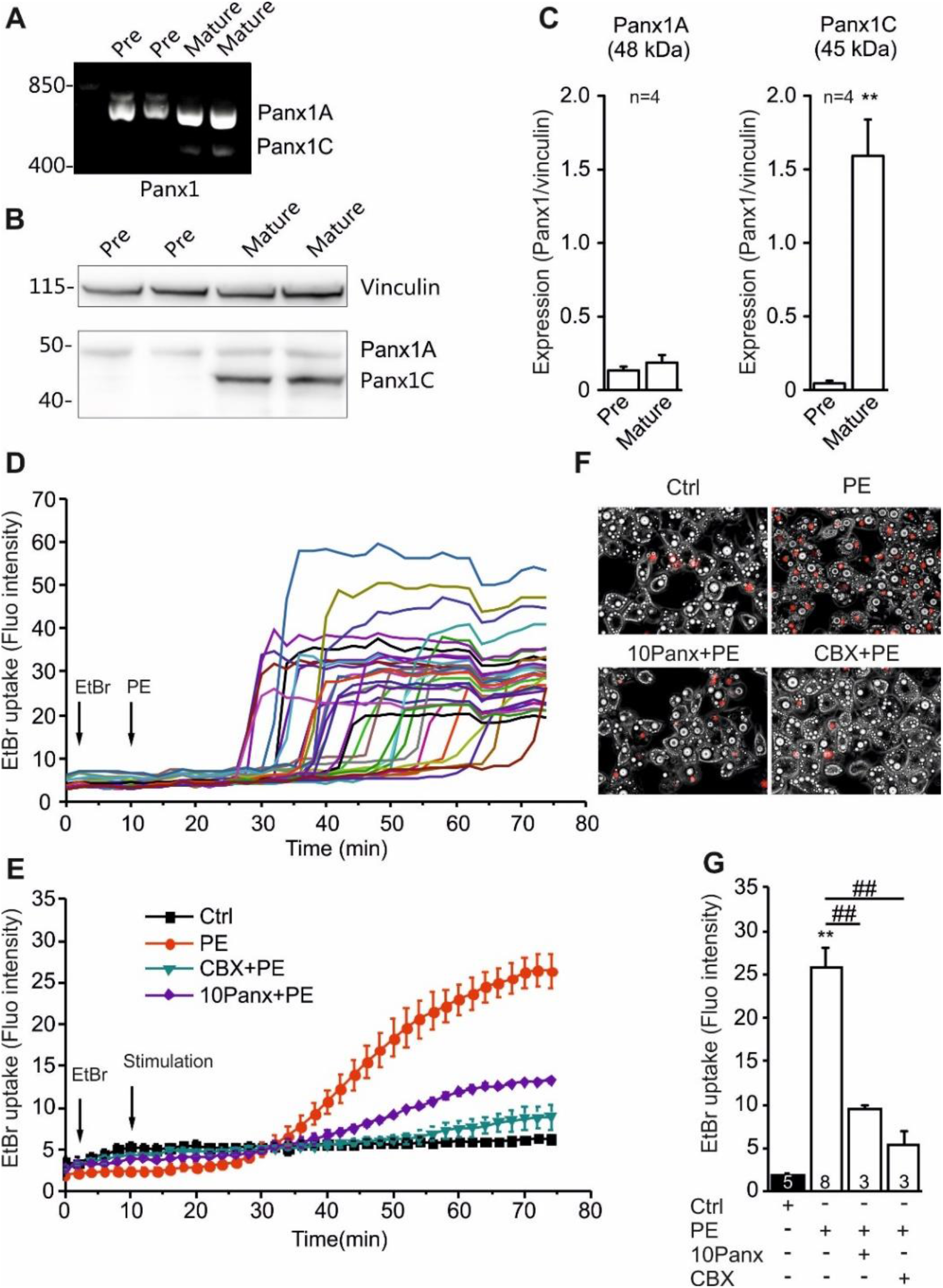
Adipocytes express Panx1 that is involved in the adrenergically stimulated pore formation. **(A)** Representative gel of *Panx1* mRNA expression and **(B)** representative western blot of Panx1 protein expression in CH3H10T1/2 pre-adipocytes and mature adipocytes. Vinculin was used as a loading control. **(C)** Western blot quantification of Panx1A and Panx1C isoforms expression. **(D)** Representative recordings of EtBr (0.5 μM) uptake in single mature adipocytes (n=30) stimulated with 5 μM phenylephrine (PE) and fluorescence was recorded every 2 min for 75 min. **(E)** Representative images of brightfield (grey) and EtBr fluorescence (red) after 70 min of stimulation with control buffer (Ctrl) or phenylephrine (PE) in presence or absence of 100 μM ^10^Panx (10Panx) or 30 μM Carbenoxolone (CBX). (F) The time-course of EtBr uptake after stimulation with control buffer (Ctrl) or phenylephrine (PE) alone, and in combination with 100 μM ^10^Panx or 30 μM CBX. The graph shows the mean ± s.e.m. responses of 90 cells per condition per independent experiment. (g) Quantification of EtBr uptake as difference between the basal fluorescence (before stimulation) and the peak uptake (70 min) with the indicated treatment. All data are shown as mean values ± s.e.m. and significance related to the control condition is indicated p<0.05 (*) and p<0.01 (**), while significance among conditions is indicated p<0.05 (#) and p<0.01 (##)

In order to further assess Panx 1 activity as a “pore” (27,28), we monitored Ethidium Bromide (EtBr) uptake after adrenergic stimulation. Time course measurements of single-cell fluorescence shows that phenylephrine induced step-like EtBr uptake in individual cells, but the cell response was not fully synchronized (Fig. 2D). Nevertheless, averaging of responses of a cell population in several experiments yielded a curve that shows a gradual increase of fluorescence reaching maximum at 50-70 min after stimulation (Fig. 2E). Pore formation was tested in experiments where phenylephrine was given alone, and in combination with ^10^Panx or carbenoxolone (Fig. 2E-G). Importantly, the phenylephrine-induced EtBr uptake was dramatically diminished by both Panx1 inhibitors (Fig. 2E-G). This indicates that adrenergically induced dye uptake is Panx1 dependent and consistent with the ATP release.

**Fig. 3.**
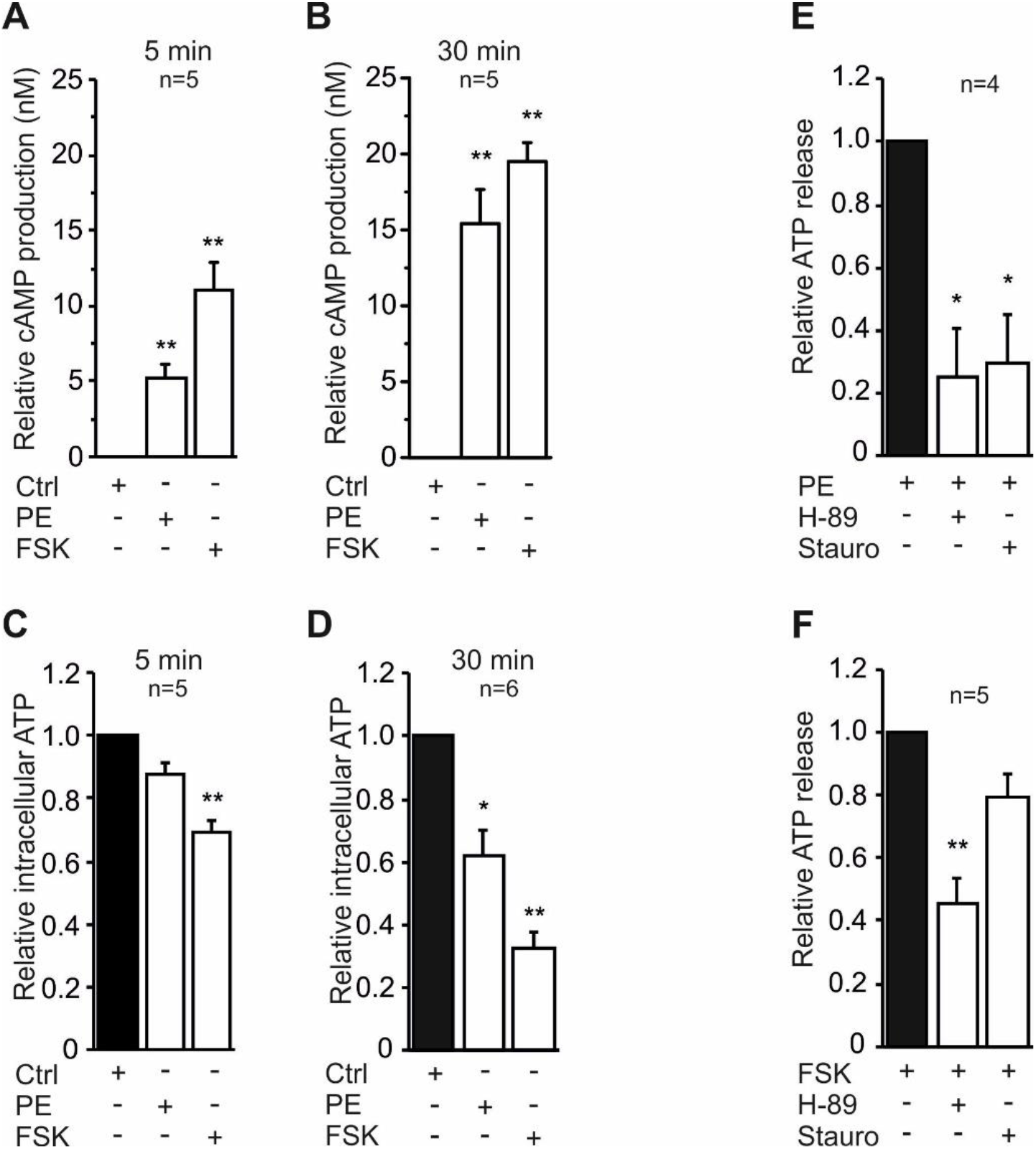
Adrenergically stimulated intracellular signaling influences ATP release. **(A, B)** cAMP production after 5 min (a) and 30 min (b) stimulation with the control buffer (Ctrl), 5 μM phenylephrine (PE) or 10 μM forskolin (FSK). **(C, D)** Normalized intracellular ATP levels after 5 min (c) and 30 min (d) stimulation with the control buffer (Ctrl), 5 μM phenylephrine (PE) or 10 μM forskolin (FSK). **(E, F)** Effect of 20 μM H-89 (PKA inhibitor) and 10 nM Staurosporine (PKC inhibitor) on phenylephrine-(e) and forskolin-(f) induced ATP release. All data are shown as mean values ± s.e.m., n indicates the number of experiments, and significance is indicated p<0.05 (*) and p<0.01 (**).

### ATP release is regulated by cAMP and PKA pathway

Next, we aimed to define the signaling pathway mediating the adrenergically induced ATP release in mature adipocyte. Since forskolin, but not ionomycin, stimulated ATP release, we focused on cAMP signaling. We measured cAMP production in mature adipocytes stimulated with phenylephrine and forskolin. At 5 and 30 min, both agonists significantly increased cAMP levels (Fig. 3A, B). In parallel experiments, both phenylephrine and forskolin reduced intracellular ATP levels compared to the control both at 5 and 30 min (Fig. 3C, D).

Having established that cAMP was significantly elevated in stimulated cells, we assessed the possible involvement of PKA, and also PKC, in the signaling pathway. Inhibition of PKA with the selective inhibitor H-89 impaired both phenylephrine- and forskolin-stimulated ATP release (Fig. 3E, F), while inhibition of PKC by staurosporine significantly decreased only the phenylephrine induced ATP release. Inhibition of exchange protein directly activated by cAMP (EPAC) did not have any effect on forskolin stimulated ATP release (data not shown).

### ATP release is coordinated with cell metabolism

To explore whether ATP release was linked to cell metabolism, we measured oxygen consumption rate (OCR) after phenylephrine and forskolin stimulation. Both agonists increased OCR in adipocytes within a few minutes and OCR continued to increase, approaching a plateau after 30 to 60 min, depending on the agonist (Fig. 4A). The mitochondrial uncoupler FCCP was used as a positive control to assess the maximal rate of respiration. Quantification of the area under the curve shows that both phenylephrine and forskolin significantly stimulated OCR compared to the control (Fig. 4B). Furthermore, to test the direct relation between metabolism and ATP release, we evaluated the effect of oligomycin, an intracellular ATP synthase blocker, on ATP release. Consistent with the OCR data, oligomycin markedly diminished the release of ATP (Fig 3C, D). This indicates that intracellular ATP formation is a prerequisite for ATP release.

**Fig. 4.**
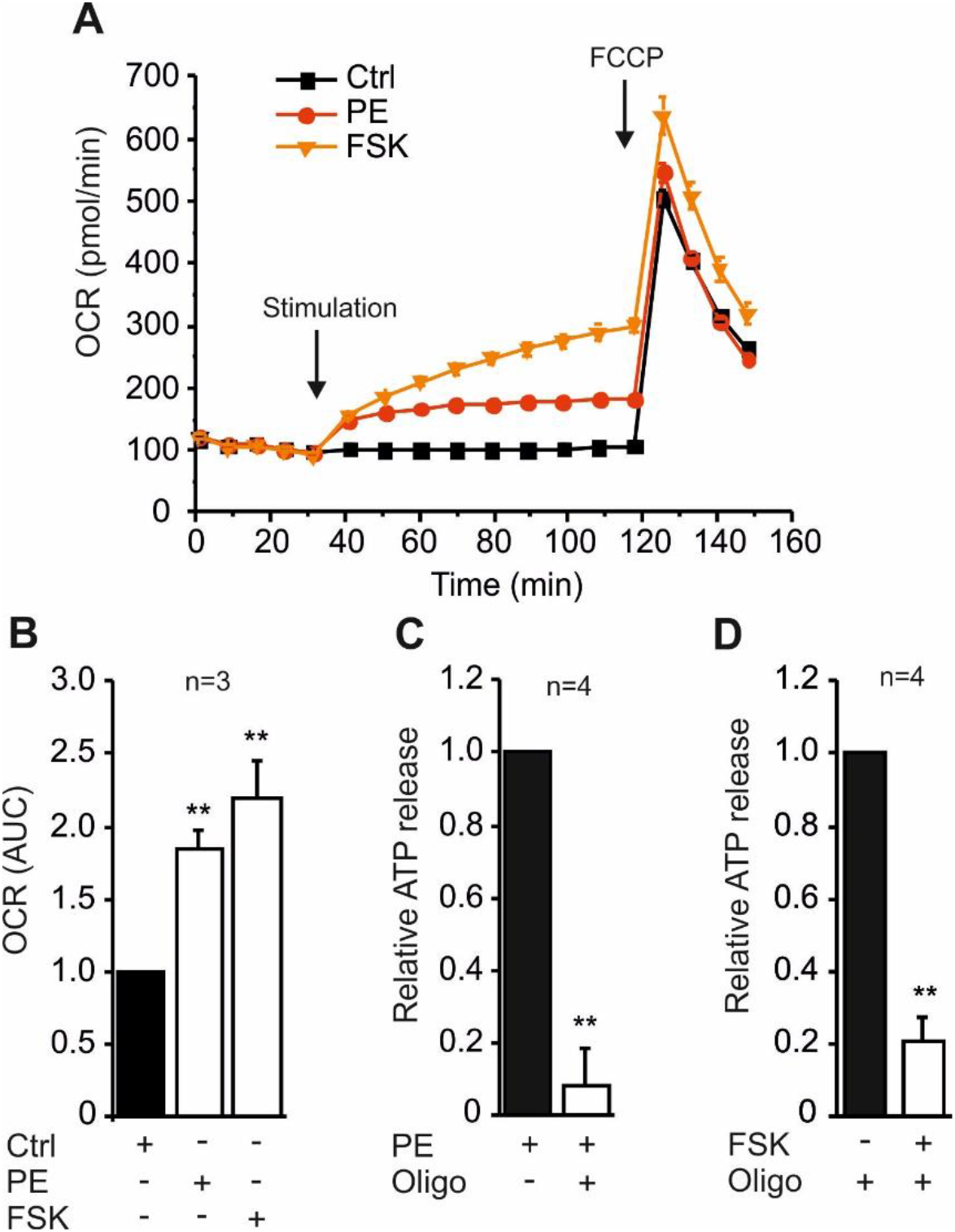
ATP release is coordinated with cell metabolism and intracellular ATP formation. **(A)** The time-course of oxygen consumption rate (OCR) under basal conditions and during successive addition of control buffer (Ctrl), 5 μM phenylephrine (PE), 10 μM forskolin (FSK), and 1 μM FCCP in a representative experiment in mature CH3H10T1/2 white adipocytes. (B) Quantification of OCR calculated as an area under the curves (AUC). **(C, D)** Effect of 1 μM oligomycin on phenylephrine-(c) and forskolin-(d) induced ATP release. All data are shown as mean values ± s.e.m., n indicates the number of experiments, and significance is indicated p<0.01 (**).

### ATP release depends on glucose and is regulated by insulin

Since cell metabolism and ATP release seemed to correlate, we investigated whether changes in extracellular glucose concentrations from 5.5 to 25 mmol/l could affect ATP release in mature adipocytes. In contrast to adrenergic stimulation, no immediate effect on extracellular ATP was observed with glucose stimulation (Fig. S1A). However, longer glucose exposure had a significant impact on adrenergically stimulated ATP release. Cells incubated in high glucose (25 mM) for 24 h and stimulated with phenylephrine, isoproterenol or forskolin markedly enhanced ATP release compared to cells incubated with the lower glucose concentration (5.5 mM) (Fig. 5A).

**Fig. 5.**
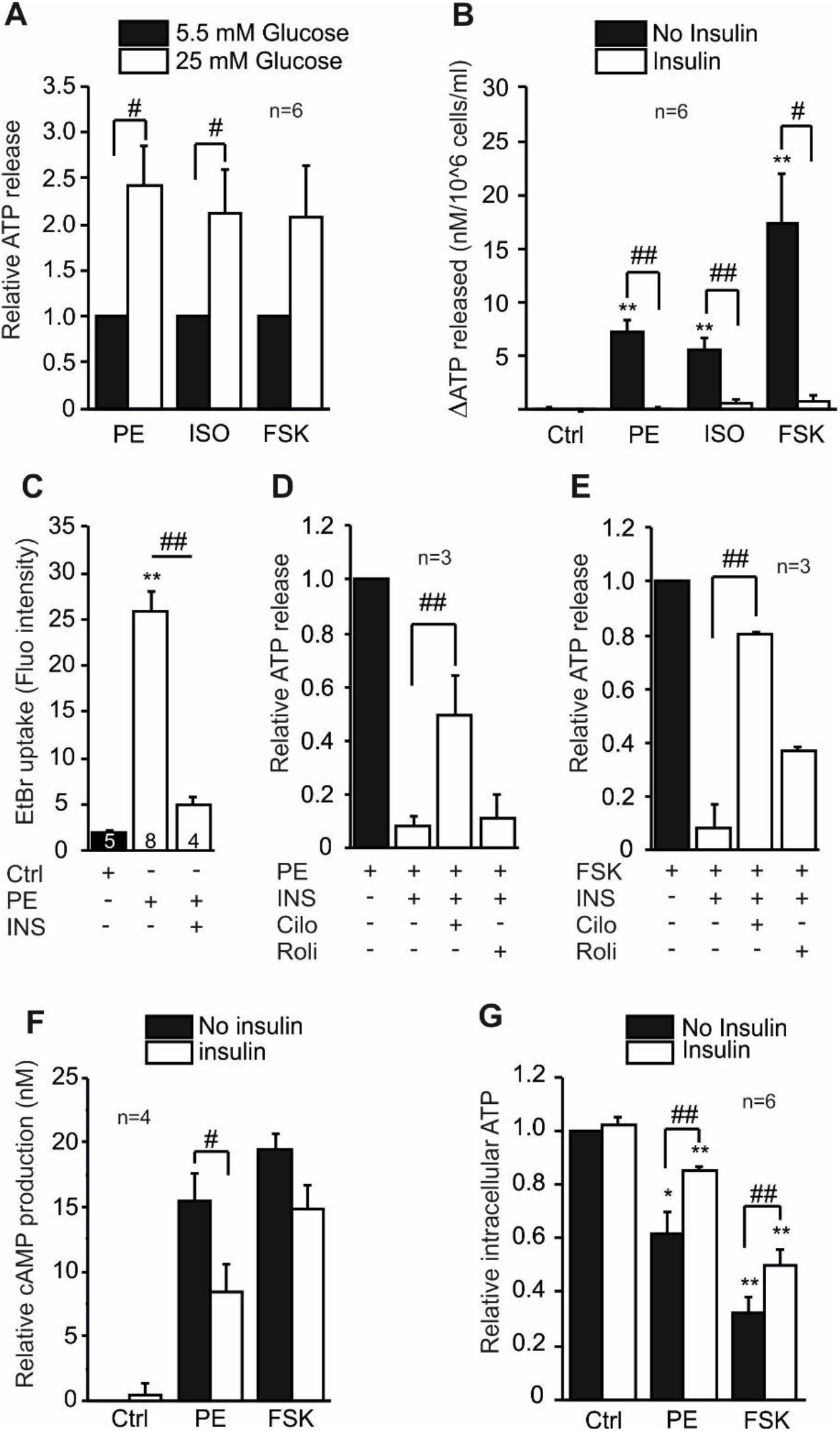
Effect of glucose and insulin on ATP release, cAMP and intracellular ATP. **(A)** Normalized ATP release in adipocytes cultured for 24 h in low (5.5 mM) or high (25 mM) glucose concentrations and then stimulated with 5 μM phenylephrine (PE), 1 μM isoproterenol (ISO) or 10 μM forskolin (FSK). **(B)** Quantification of ATP release after stimulation with the control buffer (Ctrl), 5 μM phenylephrine (PE), 1 μM isoproterenol (ISO) and 10 μM forskolin (FSK) in presence or absence of 1 μg/ml insulin. **(C)** Quantification of EtBr uptake in adipocytes stimulated with 5 μM phenylephrine (PE) in presence or absence of 1 μg/ml insulin **(D, E)** Normalized phenylephrine (d) and forskolin (e) induced ATP release in presence of insulin and either 50 μM Cilostazol (Cilo) or 20 μM Rolipram (Roli). **(F)** cAMP production after 30 min stimulation with control buffer (Ctrl), 5 μM phenylephrine (PE) or 10 μM forskolin (FSK) in presence or absence of 1 μg/ml insulin. (g) Normalized intracellular ATP levels after 30 min stimulation with control buffer (Ctrl), 5 μM phenylephrine (PE), 10 μM forskolin (FSK) in presence or absence of 1 μg/ml insulin. All data are shown as mean values ± s.e.m., n indicates the number of experiments, and significance related to the control condition is indicated p<0.05 (*) and p<0.01 (**), while significance among conditions is indicated p<0.05 (#) and p<0.01 (##).

Given the impact of glucose on ATP release, we hypothesized that insulin could also affect ATP release processes. Most notably, insulin pre-treatment abolished phenylephrine-induced ATP release and markedly inhibited the isoproterenol and forskolin induced release (Fig. 4B). In addition, insulin also inhibited phenylephrine induced Panx1-pore formation as demonstrated by fluorophore uptake (Fig. 5C). In insulin sensitive cells, the insulin receptor initiates a cascade of phosphorylation events that lead to activation of enzymes that control many steps in cell metabolism. One of the signal transduction pathways culminates with the downstream phosphorylation of cyclic nucleotide phosphodiesterases (PDEs) (29), enzymes that hydrolyze cAMP (and cGMP). Adipocytes express mainly PDE3 and PDE4, both key players in insulin-regulated lipolysis, lipogenesis, and glucose uptake (30). Therefore, we hypothesized that PDE3 and/or PDE4 could be involved in insulin-inhibition of ATP release. Indeed, inhibition of PDE3 by Cilostazol partially rescued both phenylephrine and forskolin induced ATP release from insulin inhibition (Fig. 5D, E). In comparison, inhibition of PDE4 by Rolipram did not have significant effect (Fig. 5D, E). To support the hypothesis that insulin inhibits ATP release via modulation of cAMP levels, we determined the effect of insulin on phenylephrine and forskolin induced intracellular cAMP and ATP levels (Fig. 5F, G). As predicted, in stimulated cells, insulin treatment decreased cAMP levels (Fig. 5F) and increased intracellular ATP levels (Fig. 5G). The insulin effects were specific to adrenergically induced ATP release mechanisms, because it had no effect on mechanically induced ATP release (Fig. S1B).

### Extracellular ATP promotes calcium signaling and lipolysis and attracts macrophages

Since it was demonstrated above that mature adipocytes release ATP in a regulated manner, we next probed the potential role and the importance of extracellular ATP as autocrine and paracrine signaling molecule in adipose tissue. Here, we focused on white adipocytes and the surrounding microenvironment.

Addition of extracellular ATP, a general P2 receptor activator, or 2’-3’-O-(4-benzoylbenzoyl)-ATP (BzATP), a more specific P2X7 receptor agonist, elicited similar intracellular calcium transients in mature adipocytes (Fig. 6A). White adipocytes express several types of P2X and P2Y receptors (14) and similar calcium transients have been recorded in other white adipocytes (31). Ionomycin, as expected, evoked large calcium transients (Fig. 6A), although it had no effect on ATP release (Fig. 1A, C).

**Fig. 6.**
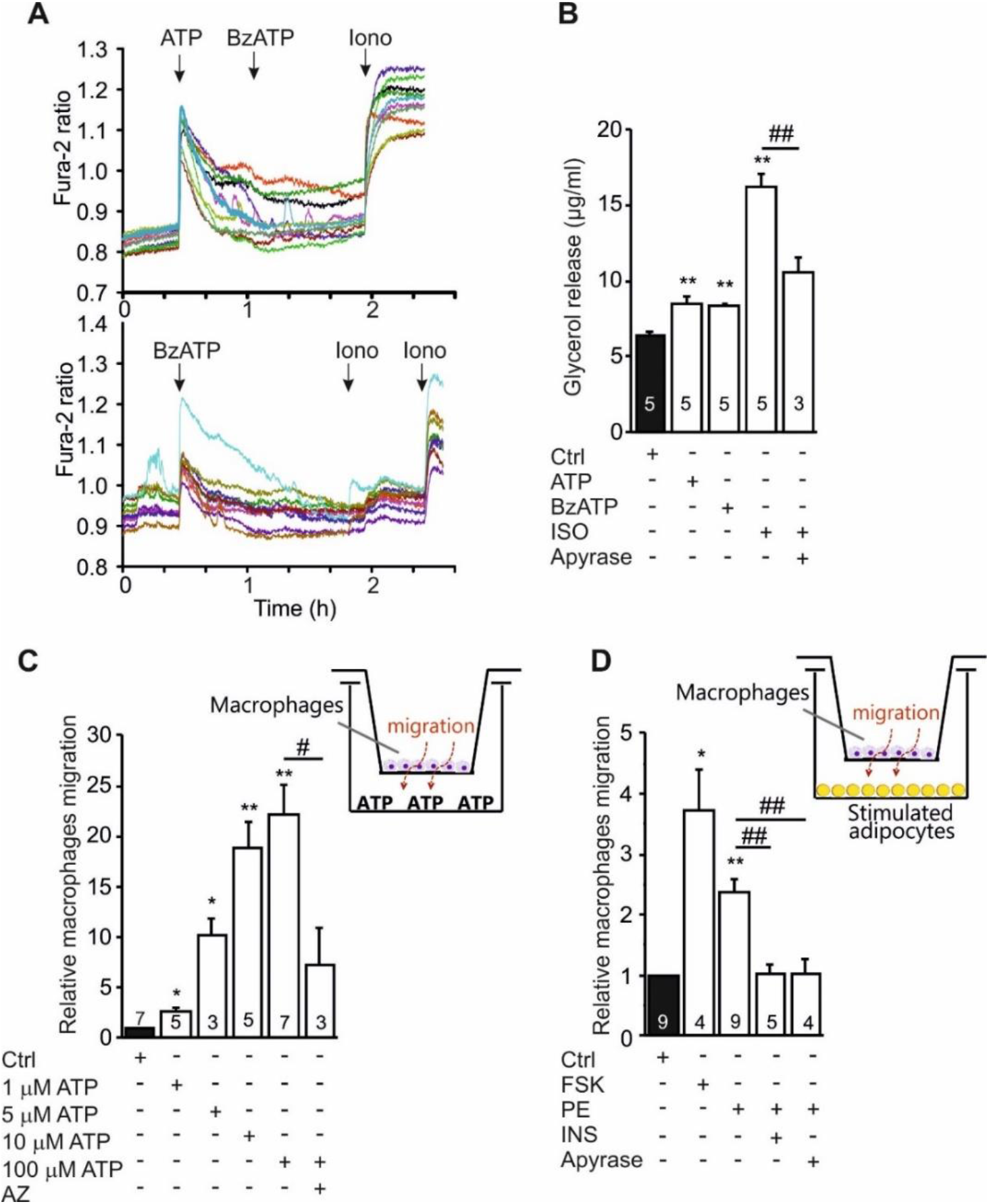
Autocrine and paracrine effect of extracellular ATP. **(A)** Representative recordings of intracellular Ca^2+^ (given as Fura-2 ratio) in white adipocytes stimulated with ATP (10 μM), BzATP (100 μM) and ionomycin (1 μM). Each figure shows responses in 10 out of 20 cells recorded per experiments, and experiments were performed 5 times. **(B)** Lipolysis in mature adipocytes treated with the control medium (Ctrl), 100 μM ATP, 100 μM BzATP and 1 μM Isoproterenol (ISO), in presence or absence of 5 U/ml apyrase. **(C)** J774A.1 macrophages migration towards increasing ATP concentrations in presence or absence of 10 μM AZ10606120 (AZ). The number of migrated cells after 3 h was normalized with respect to the control condition (Ctrl). The insert includes the schematic representation of the the Boyden chamber experimental setup. **(D)** J774A.1 macrophages migration towards ATP released by mature adipocytes stimulated with control medium (Ctrl), 10 μM forskolin (FSK), 5 μM phenylephrine (PE) alone, and in combination with 1 μg/ml insulin or 5 U/ml apyrase. The insert includes the schematic representation of the experimental setup used. All data are shown as mean values ± s.e.m. and significance related to the control condition is indicated p<0.05 (*) and p<0.01 (**), while significance among conditions is indicated p<0.05 (#) and p<0.01 (##).

Next, we tested the effect of ATP and BzATP on lipolysis. Both ATP and BzATP consistently stimulated glycerol release in C3H10T1/2 mature adipocytes (Fig. 6B). Isoproterenol was used as positive control in presence or absence of apyrase, a potent ATP-diphosphohydrolase that hydrolyses ATP to AMP. Apyrase inhibited isoproterenol-stimulated lipolysis (Fig. 6B), confirming that extracellular ATP has an important role in regulating lipolysis process in these cells. Since BzATP had significant effect, we also checked for expression of the P2X7 receptor in our adipocyte cell model and Fig. S1C, d shows that both pre-adipocytes and mature adipocytes express the P2X7 receptor.

Lastly, we investigated whether purinergic signaling mediates a cross-talk between adipocytes and macrophages. We hypothesized that higher levels of ATP released by adipocytes (in the absence of insulin inhibition) could be a chemotactic signal for immune cells, in particular macrophages. Indeed, J774A.1 macrophages migrated towards exogenous ATP (1–100 μM) in a concentration dependent manner (Fig. 6C). Interestingly, this effect seemed to be mediated at least in part by the P2X7 receptor, since the inhibitor AZ10606120 significantly reduced ATP stimulated macrophage migration (Fig. 6C).

Next, we investigated whether ATP released from adipocytes was sufficient to attract macrophages. We designed experiments (Fig. 6D) where white adipocytes were seeded and differentiated in the lower part of a Boyden chamber. Mature adipocytes were then stimulated with forskolin or phenylephrine in presence or absence of insulin and apyrase. Macrophages were seeded in the upper chamber and their migration was measured after 3 h (Fig. 6E). Clearly, macrophage migration was enhanced in experiments where adipocytes were stimulated with forskolin or phenylephrine compared to the control (Fig. 6D). Importantly, adipocytes pre-treated with apyrase or insulin did not stimulate macrophages migration (Fig. 6D), indicating that ATP release from adipocytes was the trigger for the macrophage migration.

## Discussion

In this study, we explored the mechanism and the potential role of ATP release in white adipocytes. First, we showed that adipocytes robustly released ATP in response to adrenergic stimulation. The ATP release is regulated by a cAMP-PKA dependent pathway, it is associated with increased cell metabolism, and with a particular Panx1 isoform as a release mechanism. Most importantly, we demonstrated that ATP release is enhanced by high extracellular glucose and markedly inhibited by insulin, which operates via a signaling pathway including PDE3. Furthermore, we showed that ATP released by white adipocytes is an important autocrine and paracrine regulator of adipocyte and macrophage function.

Our data show that α- and β-adrenergic receptor stimulation evokes a controlled and time dependent release of ATP in mature white adipocytes (Fig. 1A). Similarly, but a snapshot of ATP release was detected with phenylephrine in 3T3-L1 cells and isolated murine visceral adipocytes (23). Using different selective inhibitors, we showed that C3H10T1/2 adipocytes release ATP mainly through the Panx1 channel, which correlated with pore formation dynamics (Fig. 2). Perhaps surprisingly, the P2X7 receptor was not involved in the regulation of Panx1-mediated ATP release in white adipocytes (Fig. 1), although these cells express the receptor (Fig. S1C-E). In many other cells the P2X7 receptor regulates ATP efflux through Panx1 or is itself the ATP pore (6,32). A recent study showed that VNUT expression was higher in mouse and human brown adipocytes compared to white adipocytes (Razzoli et al., 2016). However, our functional study shows that the contribution of VNUT to adrenergically stimulated ATP release was modest. Interestingly, adrenergic stimulation of pre-adipocytes (undifferentiated C3H10T1/2) did not produce any release of ATP (Fig. 1B), possibly indicating that pre-adipocytes do not have the full machinery to sustain the ATP release process. Pre-adipocytes seem to express only one Panx1 isoform (Panx1A), while differentiated mature adipocytes express two isoforms, Panx1A and Panx1C, and the C isoform is markedly upregulated after differentiation (Fig. 2C). Since pre-adipocytes do not express Panx1C isoform, and they are not capable of ATP release, we propose that this novel isoform could be required for functional ATP release in mature adipocytes.

Panx1 can be activated by different mechanisms and its localization and function are also sensitive to post-translational modifications, including phosphorylation (8). In vascular smooth muscle, Panx1 contributes to ATP release and vasoconstriction mediated by the α1-adrenergic receptor (33). Mutagenesis studies indicate that Panx1 may be phosphorylated (34), however, the signaling pathway was not defined. Here, we show that adenylyl cyclase activation by forskolin induced apparently higher ATP release than adrenergic stimuli, and that Ca^2+^ ionophore had no effect. Also forskolin and phenylephrine led to cAMP accumulation and intracellular ATP depletion, consistent with the idea that both agonists, directly or indirectly, activate adenylyl cyclase, which converts ATP to cAMP, as reported for adipocytes and other cell models (35). Increased cAMP levels activate PKA, which we demonstrated to be involved in both phenylephrine and forskolin stimulated ATP release (Fig. 1E, F, Fig. 7). PKA, in turn, could directly or indirectly phosphorylate and activate Panx1 (36). In addition, PKC may also be involved in the phenylephrine-stimulated pathway (Fig. 3E), most likely modulating the indirect adrenergic activation of adenylyl cyclase (37) (Fig. 7).

**Fig. 7.**
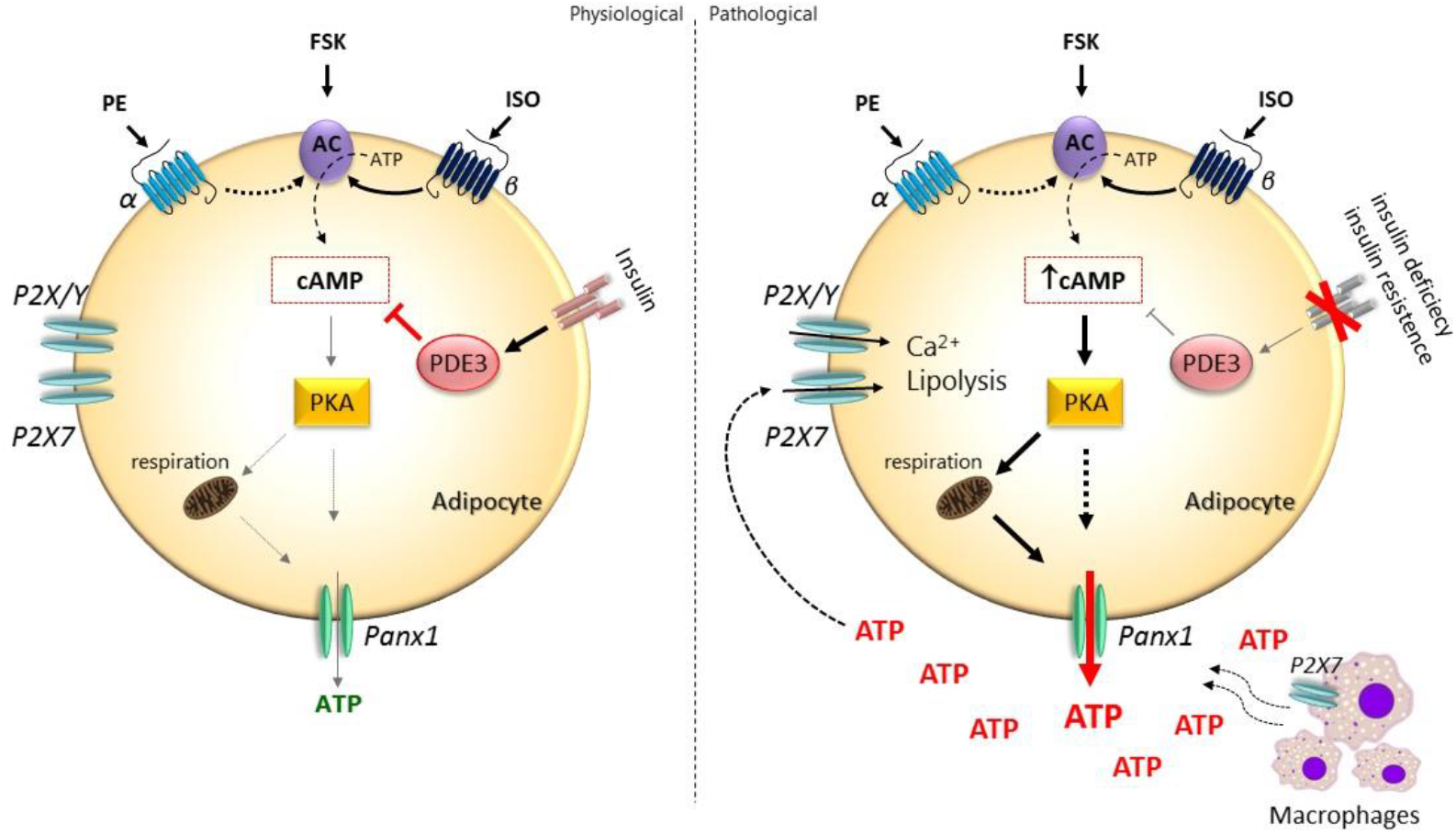
Cell model of adrenergically induced signaling resulting in ATP release and autocrine and paracrine effects in white adipose tissue. **(A)** Adrenergic stimulation of adipocytes induces cAMP increase, PKA activation, boost in cell metabolism and opening of Panx1 mediating ATP release. Insulin tightly regulates this pathway via activation of PDE3, which hydrolyses cAMP and downregulates the activation of downstream events, i.e. reduces ATP release. **(B)** In pathological conditions, such as insulin deficiency and/or insulin resistance, insulin regulation of cAMP levels would be missing, and thus PKA and Panx1 activation, and finally ATP release, would be unhindered. Extracellular ATP acts as an autocrine signal, stimulating intracellular Ca^2+^ signaling and lipolysis via P2 receptors activation. Extracellular ATP functions also as a paracrine signal, for example, as a chemoattractant for adipose tissue macrophages, stimulating their migration by activating of P2X7 receptors. The model predicts that elevated extracellular ATP (as in b) would lead to stronger and perhaps detrimental auto-/paracrine signaling in white adipose tissue.

Notably, the ATP release process in white adipocytes seems to be coordinated with increased cell metabolism and intracellular ATP production, as judged from the kinetics of these processes and oligomycin sensitivity. Indeed, the OCR elevation preceded Panx1-mediated pore formation and ATP release that was sensitive to oligomycin (Fig. 4A, C, D).

Most importantly, we demonstrate that stimulated ATP release is regulated by glucose and insulin (Fig. 5). Adipocytes grown in high glucose medium, mimicking a hyperglycaemic state, showed enhanced ATP release compared to the normoglycemic state. Remarkably, insulin markedly inhibited ATP release by stimulating the hydrolysis of cAMP via the signaling pathway that activates PDE3 (Fig. 4). This insulin-activated PDE3 signaling is known in various insulin sensitive cells, including adipocytes where it inhibits lipolysis (29,38,39). Interestingly, in erythrocytes insulin also activates PDE3 and via further regulation of CFTR (and Panx1) inhibits ATP release (40). It is not clear how insulin slightly enhanced ATP release in a suspension of visceral adipocytes, though adipocytes were not adrenergically stimulated (23).

Until now, ATP released together with norepinephrine from sympathetic nerves was considered as the main physiological source of ATP in adipose tissue. The finding that adipocytes release ATP in response to adrenergic stimulation adds an extra regulatory step that could be important in adipose tissue physiology (Fig. 7). Furthermore, the glucose and insulin sensitivity of ATP release in adipocytes may cast a new light on how this process could be deregulated during pathological states, such as hyperglycaemia, insulin deficiency or insulin resistance, revealing potential implications for type-2 diabetes. Based on our *in vitro* results we propose that in physiological conditions and under the strict control of insulin, adipocytes release very little ATP even if stimulated. In contrast, pathological conditions, such as type-2 diabetes, where cells are exposed to high glucose concentrations and the insulin control is compromised (insulin deficiency/resistance), promote ATP release in white adipocytes (Fig. 7). ATP then, via autocrine and paracrine events, affects various cell/tissue functions, and some of these we have elucidated for the white adipose tissue. This interpretation is supported by parts of previous study (23), which shows that Panx1 expression is increased in adipose tissue of obese humans and correlates with insulin resistance.

Here we show several downstream effects of ATP release. First, extracellular ATP release stimulates P2 receptors that give rise to intracellular calcium transients, which could activate various responses (14). Second, we show that ATP is a potential autocrine signal that activates lipolysis, leading to release of glycerol (Fig. 6). Notably, in our cells, ATP and BzATP have similar effect on calcium signals and lipolysis, indicating that the P2X7 receptor mediates these processes, even though we cannot exclude a role for other P2 receptors, as reported in rat white adipocytes (41). Furthermore, since apyrase inhibited part of the adrenergic effect on glycerol release, we propose that ATP release, followed by P2 receptor stimulation, potentiates the effect of adrenoceptor stimulation. Excessively elevated extracellular ATP could contribute to over-stimulation of the lipolytic process with a potentially deleterious effect on the whole-body insulin sensitivity. Sustained lipolysis and consequent release of free fatty acids (FFA) are known factors in the pathogenesis of insulin resistance, diabetes and cardiovascular diseases (10,42)

Adipose tissue includes cell types other than adipocytes, and their interaction via paracrine loops is important for adipose tissue physiology and pathophysiology (43,44). In our study, we focused on the possible cross-talk between purinergic pathways of adipocytes and macrophages, which is of particular interest with respect to adipose tissue inflammation and diabetes (45). Recent studies suggest that obesity-induced inflammation is mainly mediated by resident immune cells, especially adipose tissue macrophages (16,46). Macrophages are sensitive to ATP (47) and can be recruited by cells that release ATP (5,48,49). Our study contributes to the understanding of the underlying mechanisms. Extracellular ATP stimulated macrophage migration in a concentration- and P2X7 receptor-dependent manner (Fig. 6), a receptor expressed in macrophages (50). Most importantly, adrenergically stimulated adipocytes recruit macrophages via Panx1-mediated release of ATP in the absence of insulin. In the presence of insulin, macrophage recruitment was obliterated (Fig. 6).

In conclusion, our study revealed the importance of Panx1-mediated ATP release as a new glucose- and insulin-sensitive process in white adipocytes. We decipher the signalling pathway leading to activation of a novel splice variant of Panx1. We propose that white adipocytes release more ATP if insulin signaling is not operational and provide evidence for a role of extracellular ATP and purinergic signaling as an important autocrine and paracrine regulator of adipose tissue microenvironment. Our work provides a novel framework for understanding adipose tissue inflammation, and link between obesity and type-2 diabetes.

## Materials and Methods

### Cell culture and chemicals

We used murine the white pre-adipocyte cell line C3H10T1/2 (51) (kindly provided by Professor Karsten Kristiansen). Cells were propagated in Dulbecco’s Modified Eagle’s Medium (DMEM) (Thermofisher) supplemented with 10% fetal bovine serum (FBS), 1% penicillin/streptomycin and 25 mM glucose. One day post-confluent C3H10T1/2 cells (designated day 0) were induced to differentiate in medium supplemented with 1 μM dexamethasone (Sigma-Aldrich, D1756), 0.5 mM isobutylmethylxanthine (Sigma-Aldrich, I5879), 5 mg/ml insulin (Roche, 11376497001) and 1 μM rosiglitazone (Cayman Chemicals, 71740). At day 2, cells were refreshed with medium containing 5 mg/ml insulin and 1 μM rosiglitazone. At day 4, C3H10T1/2 cells were refreshed with medium containing 1 μM rosiglitazone. Murine macrophage J774A.1 cells (ATCC TIB-67) were propagated in DMEM supplemented with 10% fetal bovine serum (FBS), 1% penicillin/streptomycin and 25 mM glucose. All cells were cultured in a humidified atmosphere at 37°C with 5 % CO_2_. All standard chemicals were purchased from Sigma-Aldrich unless otherwise stated.

### ATP release

Extracellular ATP was monitored using luciferin/luciferase luminescence reaction in the extracellular milieu as previously described (Kowal et al., 2015). Briefly, mature adipocytes were plated (80,000 cells/well) on 96-well COSTAR white plates and grown in 5.5 or 25 mM glucose 24 h before experiment. Cells were washed and allowed to rest in physiological buffer (140 mM NaCl, 1 mM MgCl_2_, 1.5 mM CaCl_2_ 0.4 mM KH_2_PO_4_, 1.6 mM K_2_HPO_4_, 10 mM HEPES and 5.5 or 25 mM glucose, pH 7.4). After 30 min, luciferin/luciferase mix (BioThema, ATP kit SL144-041) was added and the baseline was recorded for at least 9 min. Cells were then gently stimulated with physiological buffer (control), 5 μM phenylephrine (Sigma-Aldrich, P6126), 10 μM forskolin (Sigma-Aldrich F6886), 1 μM isoproterenol (Sigma-Aldrich, I5627), 1 μM ionomycin (Sigma-Aldrich, I0634). In some experiments cells were pre-incubated 30 min with 1 μg/ml of insulin or one of the following inhibitors: sodium glyoxylate (50 μM, Sigma-Aldrich, G4502), AZ10606120 (10 μM, Tocris, 3323), ^10^Panx (100 μM, Tocris, 3348/1), Carbenoxolone (30 μM, Sigma-Aldrich, C4790), H-89 (20 μM, Calbiochem, 371963), Staurosporine (10 nM, Sigma-Aldrich, J4400), oligomycin (1 μM, Sigma-Aldrich, 75351), Cilostazol (50 μM, Sigma-Aldrich, C0737) or Rolipram (20 μM, Sigma-Aldrich, R6520). Measurements were performed at 37°C in FLUOstar Optima (BMG, Labtech). All measurements were run in triplicates. For each experimental protocol a standard curve was made with ATP (ATP standard from BioThema) concentrations ranging from 10^−10^ to 10^−5^ M. The effect of all applied stimuli and inhibitors on luciferin/luciferase assay activity was tested independently on standard curves and no effects were observed (in agreement with our earlier study (52)). Cell numbers were determined by cell counting kit-8 (CCK-8, Sigma-Aldrich, 96992) in parallel wells with same cell seeding, and the concentration of extracellular ATP was corrected to 10^6^ cells per 1 ml. The results in the bar graphs are presented as the difference between the basal concentration of extracellular ATP and the peak concentration of ATP after stimulation (ΔATP released).

### Ethidium Bromide uptake

C3H10T1/2 cells were fully differentiated in 3 cm petri dishes. AT day 6, cells were washed and allowed to rest 30 min in 900 μl of bicarbonate-containing physiological buffer (120 mM NaCl, 25 mM NaHCO_3_, 1 mM MgCl_2_ 1.5 mM CaCl_2_ 0.4 mM KH_2_PO_4_, 1.6 mM K_2_HPO_4_, 10 mM HEPES and 5.5 mM glucose, pH 7.4) in presence or absence of 1 μg/ml of insulin, 100 μM ^10^Panx or 30 μM Carbenoxolone. After the resting period, time lapse experiments in the Nikon BioStation IM-Q were started, taking images every 2 min in phase contrast and red fluorescence (Ex. 480 nm/Em. 620 nm) in 20x objective. After 2 min cells were incubated with 500 nM Ethidium Bromide and 8 min after cells were stimulated with 5 μM phenylephrine. Cells were kept in a temperature- and gas-controlled environment (37°C with 5 % CO_2_/air). Images were analyzed with Nikon NIS-Elements AR 4.13.03 software, and fluorescence intensity of at least 90 single cells per experiment was quantified.

### RNA isolation and RT-PCR

C3H10T1/2 pre-adipocytes and mature adipocytes were cultured normally, and total RNA was extracted using RNeasy Mini Kit (Qiagen) according to the manufacturer’s instructions. Primers used were as follows: Panx1 (698 + 477 bp), 5 -GAC TGG AGC TGG CGG TGG ACA AGA T-3’ (forward), 5 -GCG ATC GGG GAT GGT GCT GTC ATT T-3’ (reverse) (53); and P2X7 (128 bp), 5-TGG ATG ACA AGA ACA CGG ATG-3’ (forward), 5-CAG GAT GTC AAA ACG GAT GC-3’ (reverse) (54). 0.5 μg RNA per reaction was used in OneStep RT-PCR Kit (Qiagen) with amplification parameters as follows: one cycle at 50°C for 30 min and one cycle at 95°C for 15 min followed by 37 cycles at 94°C for 1 min, 59.3 (Panx1) or 50.4°C (P2X7R) for 1 min, 72°C for 1 min, and finally, one cycle at 72 °C for 10 min. All transcripts were run electrophoretically on 0.8% (Panx1) or 1.2% (P2X7R) agarose gels.

### Western blot

Proteins were extracted from C3H10T1/2 pre-adipocytes and mature adipocytes and samples were reduced by 10 min heating at 98°C in presence of 70 μM dithiothreitol. Proteins (30 μg) were loaded on 10 % polyacrylamide precast gels (Invitrogen) and blotted onto PVDF membranes (Invitrogen). Membranes were blocked with 5% skim milk solution in TBS-Tween (0.1 %) buffer for 1 h at room temperature and incubated overnight at 4°C with primary antibody against P2X7R (1:500 rabbit polyclonal, Alomone APR-004), pannexin-1 (1:1000 rabbit polyclonal, Alomone ACC-234) and vinculin (1:1000 mouse monoclonal, Sigma-Aldrich V9131). The blots were incubated with appropriate secondary HRP-conjugated antibodies (1:2500, Dako or Santa Cruz Biotechnology) and developed with EZ-ECL (Biological Industries) and visualized on Fusion FX (Vilber Lourmat). Band intensity was calculated using Bio1D software.

### Intracellular ATP measurements

C3H10T1/2 mature adipocytes (day 6) were plated (80,000 cells/well) on 96-well COSTAR white plate. After 24 h, cells were washed and incubated 30 min in physiological buffer containing 5.5 mM glucose in presence or absence of 1 μg/ml insulin. Cells were then stimulated 30 min with physiological buffer, 5 μM phenylephrine, 10 μM forskolin. Next, incubation buffer was removed, cells were permeabilized with digitonin (50 μM for 5 min, shaker) and the amount of intracellular ATP was assessed using luciferin/luciferase luminescence reaction (see above).

### cAMP measurements

C3H10T1/2 mature adipocytes (day 6) were plated (80,000 cells/well) on 96-well COSTAR white plate. After 24 h, cells were washed and incubated 30 min in physiological buffer containing 5.5 mM glucose in presence or absence of 1 μg/ml insulin. Cells were then stimulated 30 min with physiological buffer, 5 μM phenylephrine or 10 μM forskolin. Intracellular cAMP was measured by cAMP-Glo assay kit (Promega, V1501) according to manufacturer’s instruction.

### Seahorse measurements

Fully differentiated C3H10T1/2 cells (day 6) were plated (40,000 cells/well) on XF96 cell culture microplate, and real-time measurements of OCR (oxygen consumption rate) was performed using the Seahorse XF96 Extracellular Flux Analyzer (Seahorse Bioscience). One hour before the first measurement, the cell culture medium was changed to DMEM (Seahorse Bioscience) supplemented with 2% BSA and 5 mM glucose, pH 7.4. OCR was measured under basal conditions and during successive addition of 5 μM phenylephrine or 10 μM forskolin and 1 mM FCCP (Carbonyl cyanide 4-(trifluoromethoxy) phenylhydrazone, Sigma-Aldrich C2920).

### Calcium signals

C3H10T1/2 cells were grown and differentiated in glass bottom dishes (Willco, KIT-3512). Mature adipocytes were loaded with Fura-2 AM (3 μM) for 30 min in physiological buffer, gently washed and mounted in temperature regulated chamber (37°C). Experiments were conducted in standing-bath configuration to avoid mechanical disturbance. Cells were stimulated with ATP or 2’-3’-O-(4-benzoylbenzoyl)-ATP (BzATP, Sigma-Aldrich, B6396) and 1 μM ionomycin. Changes in intracellular Ca^2+^ were followed in Nikon Eclipse Ti microscope with 40x NA1.4 objective. Fura-2 – loaded cells were illuminated at λ_ex_ = 340 nm for 60 ms and λ_ex_ = 380 nm for 60 ms at 1 s intervals using a TILL Polychrome monochromator. Emission was collected at 500 nm by an image-intensifying, charge – coupled device (CCD) camera (Andor X3 897) and digitized by FEI image processing system (Thermo Fischer Scientific). LA Live Acquisition software was used to both control the monochromator and the CCD camera. The intracellular Ca^2+^ was presented as the ratio of Fura-2 fluorescence signals recorded at 340/380 nm.

### Glycerol release

C3H10T1/2 mature adipocytes (day 6) were plated (80,000 cells/well) on 96-well plate. After 24 h, cells were washed and incubated 4 h in serum-free buffer (DMEM with 2% fatty acid-free BSA (Sigma-Aldrich A8806) and 25 mM glucose). Next, cells were stimulated with 100 μM ATP, 100 μM BzATP or 1 μM Isoproterenol. Where indicated, cells were pre-incubated with 5 U/ml apyrase (Sigma-Aldrich, A6132) for 30 min prior stimulation. Cell culture medium was then collected after 6 h of stimulation and stored at −20°C. Glycerol release was measured according to the manufacturer’s instructions using the Adipolysis Assay Kit (Cayman Chemical, 10009381).

### Migration assay

J774A.1 macrophages were washed and resuspended in serum-free buffer (DMEM with 0.1% fatty acid-free BSA and 25 mM glucose). 10^6^ cells/ml were seeded in the upper chamber of Boyden Chambers (transparent PET membrane, 8.0 μm pore size, Falcon). ATP was diluted to the indicated concentrations in serum-free buffer and added into the lower chamber. Alternatively, C3H10T1/2 cells were seeded and differentiated in the lower chamber. At day 6, adipocytes were washed and incubated for 30 min in serum-free buffer in presence or absence of 1 μg/ml of insulin or 5 U/ml apyrase. Next, adipocytes were simulated with serum-free buffer, 10 μM forskolin or 5 μM phenylephrine, following by immediate addition of macrophages into the upper chamber. After 3 h incubation at 37°C in a humidified atmosphere with 5% CO_2_/air, migrated J774A.1 macrophages were fixed in cold methanol and stained with Crystal Violet. Bright field images of 8 fields of view per insert were taken with 10x objective in Leica DMI6000B microscope. Cells were counted by ImageJ, NIH. The effect of forskolin, phenylephrine, insulin and apyrase on macrophages migration was also tested in absence of adipocytes and used to normalize the corresponding treatment in presence of adipocytes.

### Statistics

Data are shown as mean ± standard error of the mean (s.e.m.) and n denotes the number of independent experiments. Sigma Plot 13 was used for the following analyses. Data were tested for normality. Normalized data were analyzed with One-sample t-tests, followed by correction for multiple comparisons with the Holm-Bonferroni method, when more than two different conditions were tested relative to the control. Non-normalized data were tested with Student’s t tests or One-way ANOVA followed by post-hoc Bonferroni tests. Significant differences p<0.05 (*) and p<0.01 (**) are indicated.

## Supplementary Materials

**Figure S1.**
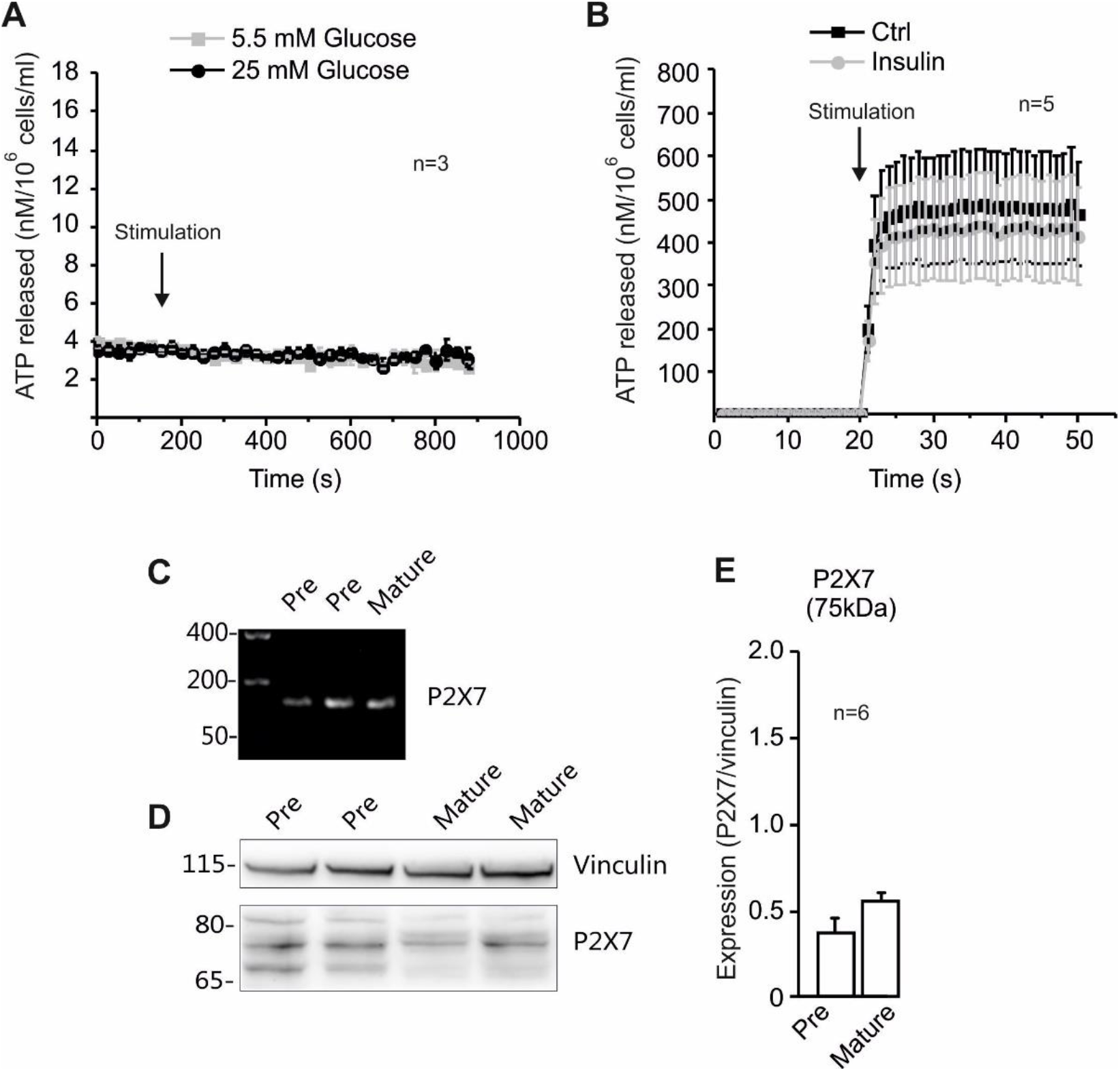
Effect of glucose and mechanical stimulation on ATP release and P2X7 receptor expression in mature white adipocytes. **(A)** The ATP release in C3H10T1/2 mature adipocytes is not affected by acute glucose load; i.e. control buffer containing 5.5 mM glucose or 25 mM glucose were given as indicated by the arrow. **(B)** The time-course of ATP release after pump injection of a control buffer at 260 μl/sec (mechanical stimulation) in presence or absence of 1 μg/ml insulin. **(C)** Representative gel of *P2X7* mRNA expression in pre- and mature C3H10T1/2 adipocytes. **(D)** Representative western blot of P2X7 protein expression. Vinculin was used as loading control. **(e)** Western blot quantification of P2X7 full length (75 kDa) expression.

## Acknowledgements

We thank L.K. Markussen for advice on adipocyte culture and for helpful discussions. We are grateful to N. M. Christensen for setting up the pore assay. Images were taken at the Center for Advanced Bioimaging, University of Copenhagen.

## Author contributions

M.T. and I.N. designed and performed the experiments and analyzed the data. M.T. and I.N. wrote the manuscript. J.B.H. and I.N. supervised the research. All authors read and approved the manuscript.

## Data and material availability

All data required to evaluate the conclusions reached in the paper are present in the paper and the paper or the Supplementary Materials.

## Supplementary material

Supplementary Fig. 1

